# TDP-43 nuclear loss in FTD/ALS causes widespread alternative polyadenylation changes

**DOI:** 10.1101/2024.01.22.575730

**Authors:** Yi Zeng, Anastasiia Lovchykova, Tetsuya Akiyama, Chang Liu, Caiwei Guo, Vidhya Maheswari Jawahar, Odilia Sianto, Anna Calliari, Mercedes Prudencio, Dennis W. Dickson, Leonard Petrucelli, Aaron D. Gitler

## Abstract

In frontotemporal dementia and amyotrophic lateral sclerosis, the RNA-binding protein TDP-43 is depleted from the nucleus. TDP-43 loss leads to cryptic exon inclusion but a role in other RNA processing events remains unresolved. Here, we show that loss of TDP-43 causes widespread changes in alternative polyadenylation, impacting expression of disease-relevant genes (e.g., *ELP1, NEFL,* and *TMEM106B*) and providing evidence that alternative polyadenylation is a new facet of TDP-43 pathology.

## Main

TDP-43 binds to uridine/guanine (UG)-rich motifs in RNA transcripts^1,2^ and plays a critical role in various aspects of RNA metabolism^3–5^. Defects in RNA metabolism are considered central to frontotemporal dementia (FTD) and amyotrophic lateral sclerosis (ALS) pathogenesis^3^. However, it is not yet fully understood what genes TDP-43 regulates, what aspect of their RNA metabolism TDP-43 regulates, and how their dysregulation promotes neurodegeneration. A major role of TDP-43 has emerged as a repressor of so-called cryptic exons during splicing^6^. Cryptic exons reside in introns of genes and are normally excluded from mature mRNAs.

When TDP-43 is dysfunctional (i.e., when depleted from the nucleus in FTD/ALS), these cryptic exons are spliced into final mRNAs, often leading to frameshifts, decreased RNA stability, or even the production of novel peptide sequences^7–13^. Importantly, some of these cryptic exons are in genes critical for neuronal functions (e.g., *STMN2*^7–9,14^) or genes that harbor disease-associated variants that sensitize them to cryptic splicing upon loss of TDP-43 (e.g., *UNC13A*^10,11^). Cryptic splicing events could serve as powerful biomarkers for TDP-43 dysfunction^12,13^ or even as therapeutic targets^15,16^.

Besides its well-established role in splicing, TDP-43 also plays a critical role in other aspects of RNA processing. Do some of these additional TDP-43-dependent RNA processing pathways also contribute to disease? One potential disease-relevant RNA processing pathway is alternative polyadenylation (APA) – a major layer of gene regulation that occurs in >60% of human genes^17^. When a gene is transcribed into mRNA, it is cleaved and polyadenylated, which functions to stabilize the mRNA, facilitate its nuclear export, and regulate its translation^18^. If alternative polyadenylation (polyA) sites are utilized, it could produce mRNA isoforms that have different 3’ untranslated region (UTR) lengths, impacting RNA/protein levels, subcellular localization, or even protein functions^17,19^. If APA occurs prematurely, it can truncate mRNAs, reducing full-length protein levels^17,19^. Previous genome-wide TDP-43 binding studies revealed that TDP-43 binding is enriched not only in introns of genes but also in 3’ UTRs^1,2^, suggesting a role in regulating polyadenylation. Indeed, TDP-43 knockdown or disease-associated TDP-43 mutations affect polyadenylation^20–22^, including its own polyadenylation^2,23^ and premature polyadenylation in *STMN2*^7,8^. Notably, widespread APA changes have been observed in ALS patient samples using bulk RNA-sequencing (RNA-seq)^24^ and single nucleus RNA-seq^25^, although it is unclear whether these changes are directly owing to TDP-43 dysfunction.

To test the hypothesis that APA caused by TDP-43 dysfunction contributes to FTD/ALS, we first performed APA analysis in an RNA-seq dataset^26^, in which neuronal nuclei with and without TDP-43 were sorted from FTD/ALS postmortem brain samples for RNA-seq (**Fig. 1a, top panel**). We recently re-analyzed this dataset to discover TDP-43-dependent cryptic splicing events^11^. Here, we ‘re-re-analyzed’ it to search for APA events (**Fig. 1a, bottom panel**). We used two different APA analysis programs and identified 41 APA changes using APAlyzer^27^ and 62 APA changes using QAPA^28^ (**Fig. 1b**, **S1a; Table S1;** adjusted *p* value < 0.1). Only two genes with significant APA changes were common between both programs: *LRFN1* and *MARK3* (see also **Fig. S2k** and Arnold et al.^29^), likely because of different polyA databases used by each program and the low resolution of RNA-seq for APA analysis. We thus considered APA changes identified by either program, which revealed APA events in genes critical for neuronal function, such as *LRFN1* and *SYN2*. Some of these APA events are associated with gene expression changes (**Fig. S1b**), likely impacting their corresponding protein levels (**see below**).

**Figure 1.**
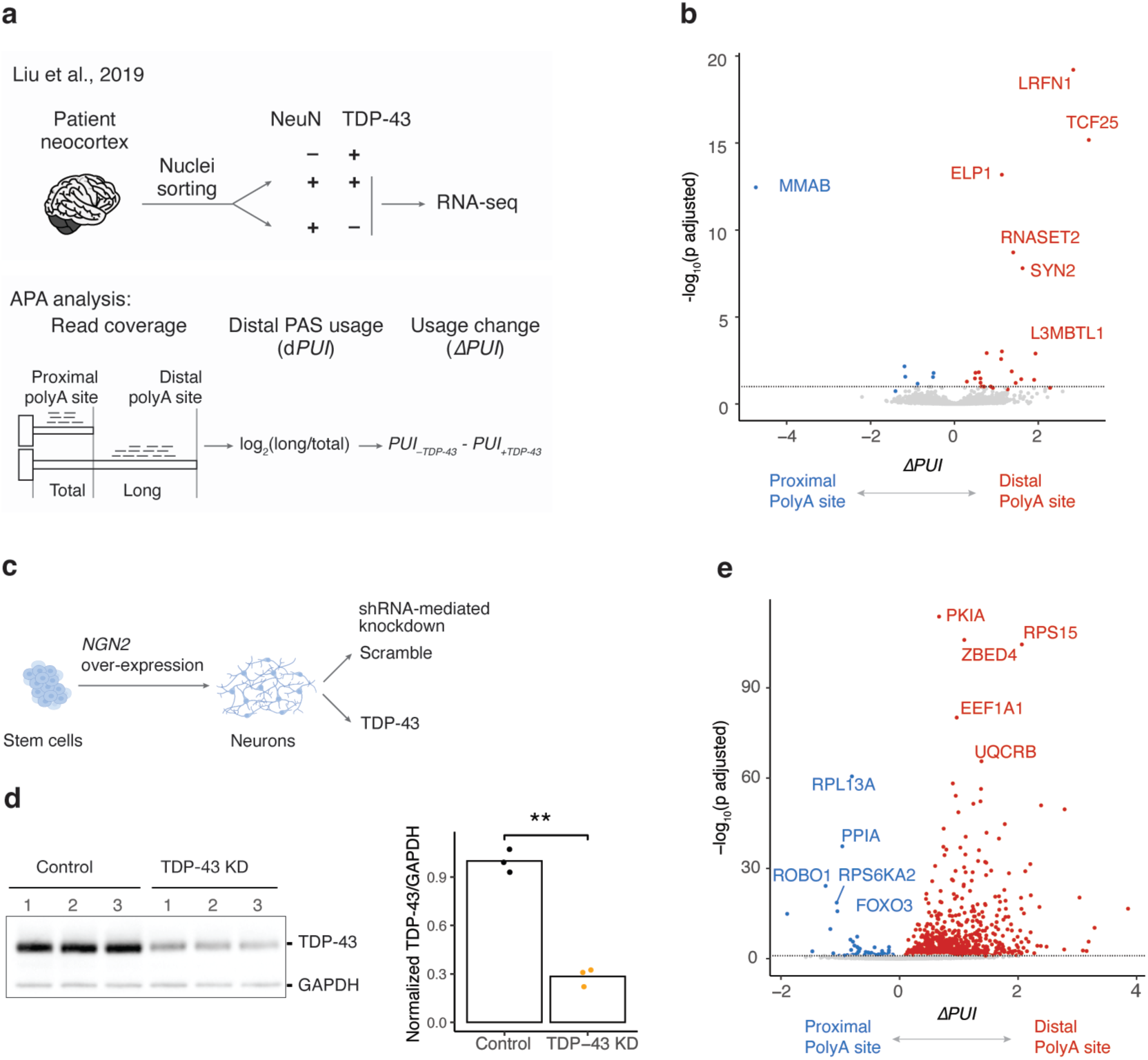
Loss of TDP-43 leads to widespread alternative polyadenylation changes. **a**, APA analysis scheme. **Top panel**, scheme for sorting neuronal nuclei from frontal cortices of FTD or FTD-ALS patients based on TDP-43 levels for RNA-seq^26^. **Bottom panel**, workflow of APA analysis using RNA-seq data. **b**, Loss of TDP-43 is associated with APA changes. APA change (*ΔPUI* by APAlyzer) is plotted on the *x*-axis and the corresponding statistical significance (the adjusted *p* value by DEXseq) on the y-axis. Genes with statistically significant APA change (adjusted *p* < 0.1) are labeled in red for favoring distal polyA sites and in blue for proximal polyA sites. Dashed horizontal line marks the adjusted *p* value of 0.1. **c**, *NGN2* over-expression mediated differentiation of human stem cells to cortical neurons for shRNA-mediated TDP-43 knockdown (KD). **d**, Western blot (WB) shows efficient TDP-43 KD after 7 days. *p*-value was calculated by Student’s *t* test. **e**, Volcano plot shows widespread APA changes upon TDP-43 KD in iNeurons; thresholds for significant changes are *|ΔPUI|* > 0.10 and adjusted *p* < 0.05 from APAlyzer. ns (not significant), *p* > 0.05; *, *p* ≤ 0.05; **, *p* ≤ 0.01; ***, *p* ≤ 0.001; ****, *p* ≤ 0.0001.

To test if TDP-43 directly regulates APA, we knocked down TDP-43 levels in cortical neurons differentiated from human stem cells (iNeurons) and performed RNA-seq for APA analysis (**Fig. 1c-e**). TDP-43 knockdown caused widespread APA changes (**Fig. 1e; Table S2;** |*ΔPUI*| > 0.1 and adjusted *p* value < 0.05 from APAlyzer), including 24 changes also observed in FTD/ALS postmortem brain samples (**Fig. 1b**, **S1a**). We also observed APA changes in induced pluripotent stem cell (iPSC)-derived motor neurons (iMNs) upon TDP-43 knockdown^7^ (**Fig. S1c**), in iNeurons carrying a pathogenic mutation (TDP-43-K263E)^30^ (**Fig. S1d**), and in iMNs carrying a different pathogenic mutation (TDP-43-M337V)^21^ (**Fig. S1e**). Thus, TDP-43 regulates APA in neurons and TDP-43 loss of function or ALS-linked mutations result in widespread APA changes.

To comprehensively map TDP-43-dependent APA events and define how TDP-43 dysfunction contributes to neurodegeneration via APA, we sought a more sensitive assay than RNA-seq for APA analysis, because RNA-seq cannot map novel polyadenylation events, identify actual polyA sites, or capture ‘internal’ premature polyadenylation events (i.e., ones that truncate the RNA/protein). We used a specialized transcriptomic method, 3’ end-seq, to map polyA sites with single-nucleotide resolution in iNeurons with or without TDP-43 knockdown (**Fig. 2a**, **1d**). We obtained a total of 117,823 putative polyA sites with a known upstream polyA signal, of which 46,760 sites are novel (**Fig. S2a**), when compared to a compendium of polyA site annotations^31^. Consistent with previous findings^2,20^, the majority of polyA sites we identified are in 3’ UTRs (**Fig. S2b**). 3’ end-seq confirmed that loss of TDP-43 lengthened the 3’ UTR of *LRFN1* (**Fig. 2b, top panel**), which we also observed in our analysis of FTD/ALS postmortem brain samples (**Fig. 2b, top panel; Fig. 1b**). Notably, 3’ end-seq also captured premature polyadenylation in *STMN2*, which was previously identified but not apparent in standard RNA-seq data (**Fig. 2b, bottom panel**), demonstrating the power of 3’ end-seq for studying premature polyadenylation, the most detrimental form of APA since it truncates the RNA/protein.

**Figure 2.**
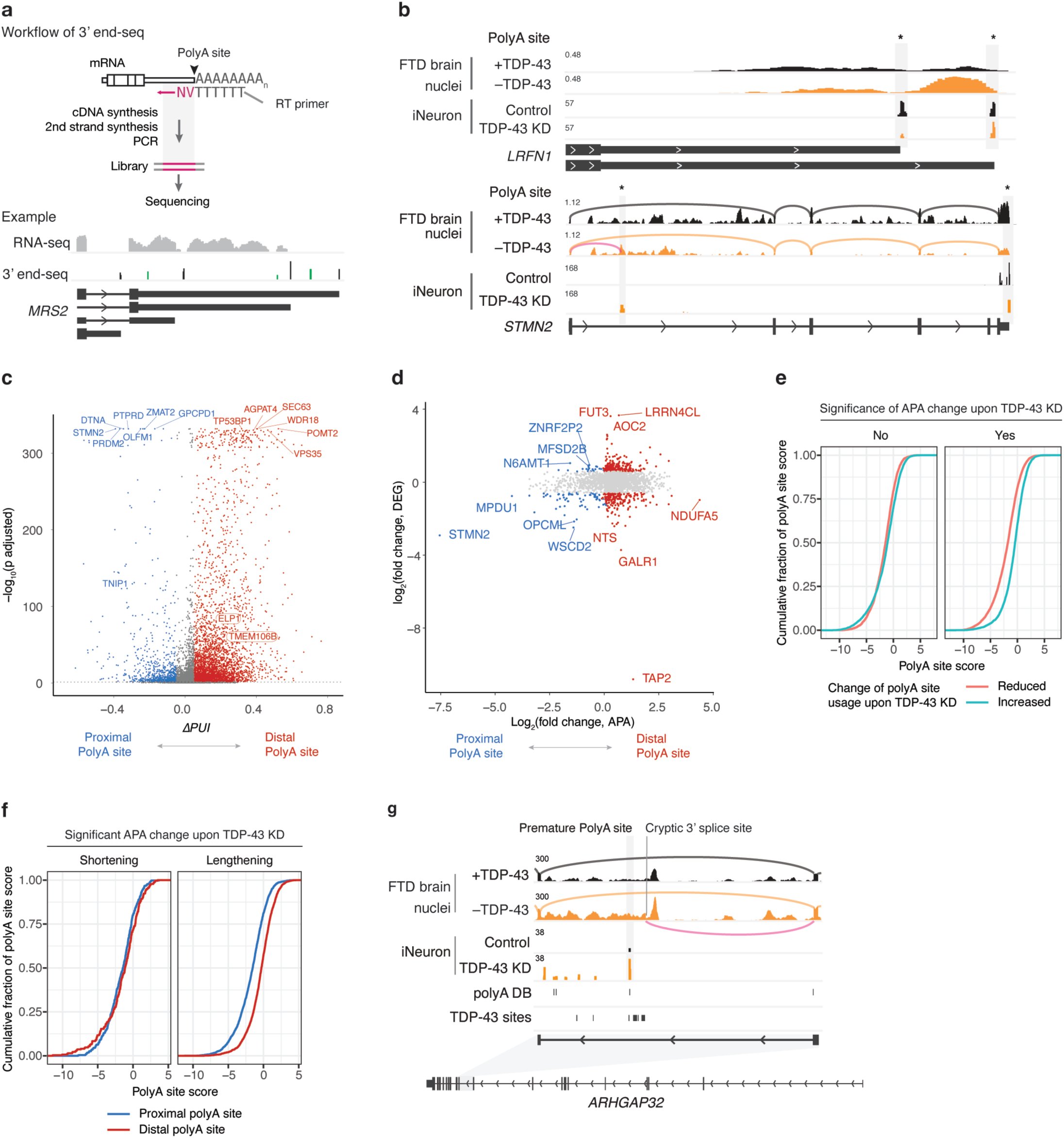
High-resolution mapping of TDP-43 dependent alternative polyadenylation using 3’ end-seq. **a**, **Top panel**, workflow of 3’ end-seq. **Bottom panel**, example of 3’ end-seq read coverage, indicative of polyA sites, at the 3’ UTR of *MRS2*. Top track: RNA-seq read coverage; middle track: read coverage of 3’ end-seq with peaks corresponding to novel sites marked in green; bottom track: gene structure. **b**, **Top panel**, example of TDP-43 regulated APA changes detected by 3’ end-seq. The direction of *LRFN1* gene is flipped for illustration purpose. **Bottom panel**, example of premature polyadenylation captured by 3’ end-seq. Sashimi plots in the top two tracks illustrate splicing; the pink curved line marks the usage of a cryptic 3’ splice site in intron 1 of *STMN2*. **c**, Volcano plot shows that 3’ end-seq captures widespread APA changes upon TDP-43 knockdown (KD). Thresholds for significant changes are *|ΔPUI|* > 0.10 and adjusted *p* < 0.05. **d**, Scatter plot with APA changes, represented by distal polyA site usage, on the *x*-axis, and RNA level changes on the *y*-axis, illustrating that APA changes could alter RNA levels. Genes with significant APA changes are plotted; genes with significant RNA level changes are labeled in red when favoring distal polyA sites or blue when favoring proximal polyA sites. **e**, Cumulative plots show that for TDP-43 regulated APA sites, distal polyA sites are stronger than proximal polyA sites. **f**, Cumulative plots show that for genes with significant 3’ UTR lengthening upon TDP-43 KD, their distal polyA sites are stronger than their proximal ones. PolyA site scores (in panels **e** and **f**) were calculated using Aparent2, expressed as log odds ratio, and reflect polyA site strengths. **g**, TDP-43 KD activated cryptic splicing and premature polyadenylation in intron 12 of *ARHGAP32*. The “polyA DB” track marks annotated polyA sites^59^ and the “TDP-43 sites” track marks the observed TDP-43 binding^32^. Gene structure is shown on the bottom.

In total, TDP-43 knockdown altered the usage of 8,169 polyA sites (|*ΔPUI*| > 0.1 and adjusted *p* value < 0.05 from LAPA, a 3’ end-seq data focused program) and caused APA changes in 2,220 genes (**Fig. 2c; Table S3**). By cross-referencing APA sites with a curated genome-wide TDP-43 binding site dataset^32^, we found that ∼70% of genes with APA events have at least one TDP-43 binding site, consistent with a direct role of TDP-43 in regulating APA of these genes. Like recent findings in ALS samples^24^ and FTD samples (**Fig. 1b**, **S1a**) as well as our RNA-seq data from iNeurons (**Fig. 1e**), the majority of APA changes (1,881) detected by 3’end-seq lengthened RNA transcripts; 340 APA events were associated with at least a 1.5-fold change in RNA level (**Fig. 2d**), suggesting that APA changes could alter gene expression. Notably, several of these APA changes were also observed in FTD/ALS postmortem brain samples and were in genes connected to ALS or FTD (**Table S4; see below**).

To define whether polyA site strength influences TDP-43 regulated APA, we estimated the probability of cleavage and polyadenylation for each identified polyA site using a deep residual neural network-based model, Aparent2^33^, that accurately predicts polyA site usage. Interestingly, we found that polyA sites with reduced usage upon TDP-43 knockdown are weaker than the ones with increased usage upon TDP-43 knockdown (**Fig. 2e**), indicating that TDP-43 either promotes the usage of weak polyA sites or suppresses the usage of strong ones. Whereas genes with significant 3’ UTR shortening show no difference in polyA strength between proximal and distal polyA sites, genes with significant 3’ UTR lengthening have stronger distal sites compared to proximal ones (**Fig. 2f**). Given the co-transcriptional nature of cleavage and polyadenylation, our findings suggest that in the case of 3’ UTR lengthening upon TDP-43 knockdown, a stronger distal polyA site might compensate for its positional disadvantage over the proximal polyA site during transcription and therefore get activated upon TDP-43 knockdown. The impact of polyA site strength on shaping TDP-43-regulated APA provides a toehold to prioritize antisense oligonucleotide-based therapeutic strategies to target specific APA events akin to current approaches underway to target cryptic splicing events^15^.

3’ end-seq also empowered the discovery of “cryptic” polyA sites (ones not used under normal conditions but revealed upon TDP-43 knockdown). TDP-43 knockdown activated 457 cryptic polyA sites in 424 genes (**Table S5**); 163 cryptic polyA sites occurred downstream of a gene’s annotated 3’ end, such as the FTD risk factor *RFNG*^34^ (**Fig. S2c**), *SIX3*, *TLX1*, and *ELK1* (**Fig. S2d-f**, and see also Bryce-smith et al.^35^) and 153 cryptic events induced premature polyadenylation in genes such as *TNIP1*, *EGFR*, *SLC24A3,* and *GSTO2* (**Fig. S2g-j**). Like coordinated activation of splicing and polyadenylation in intron 1 of *STMN2*, we observed coordinated activation in *PIGL*, *HNRNPA1,* and *ARHGAP32* (**Fig. 2g**), the latter of which is observed by Fratta and colleagues^35^. Cryptic splicing of *ARHGAP32* was recently reported^12^, and we also confirmed this in RNA-seq data from FTD/ALS postmortem brain samples (**Fig. 2g**). These results suggest a potential coupling between cryptic splicing and the activation of cryptic polyadenylation sites.

TDP-43 depletion increased the usage of two distal polyA sites in *ELP1* that lengthened its 3’ UTR (**Fig. 3a, left panel; Fig. S3a**) and such lengthening is present across different datasets (**Fig. S3b**; see also Arnold et al.^29^). *ELP1*, also called *IKBKAP,* encodes a subunit of the elongator complex, which has been functionally and genetically linked to ALS^36–38^. In addition to *ELP1*, we observed APA changes in two other subunits of the elongator complex (*ELP3* and *ELP6*) upon TDP-43 knockdown (**Fig. S3c, S3d**). These APA changes are associated with protein levels; TDP-43 knockdown led to increased protein levels of ELP1 and ELP3 (**Fig. 3a**, **S3c**). Together with a recent finding of reduced aminoacylation of tRNA^Phe^ in FTD/ALS^39^, our observations suggest that altered tRNA metabolism might be an important mechanism contributing to FTD/ALS.

**Figure 3.**
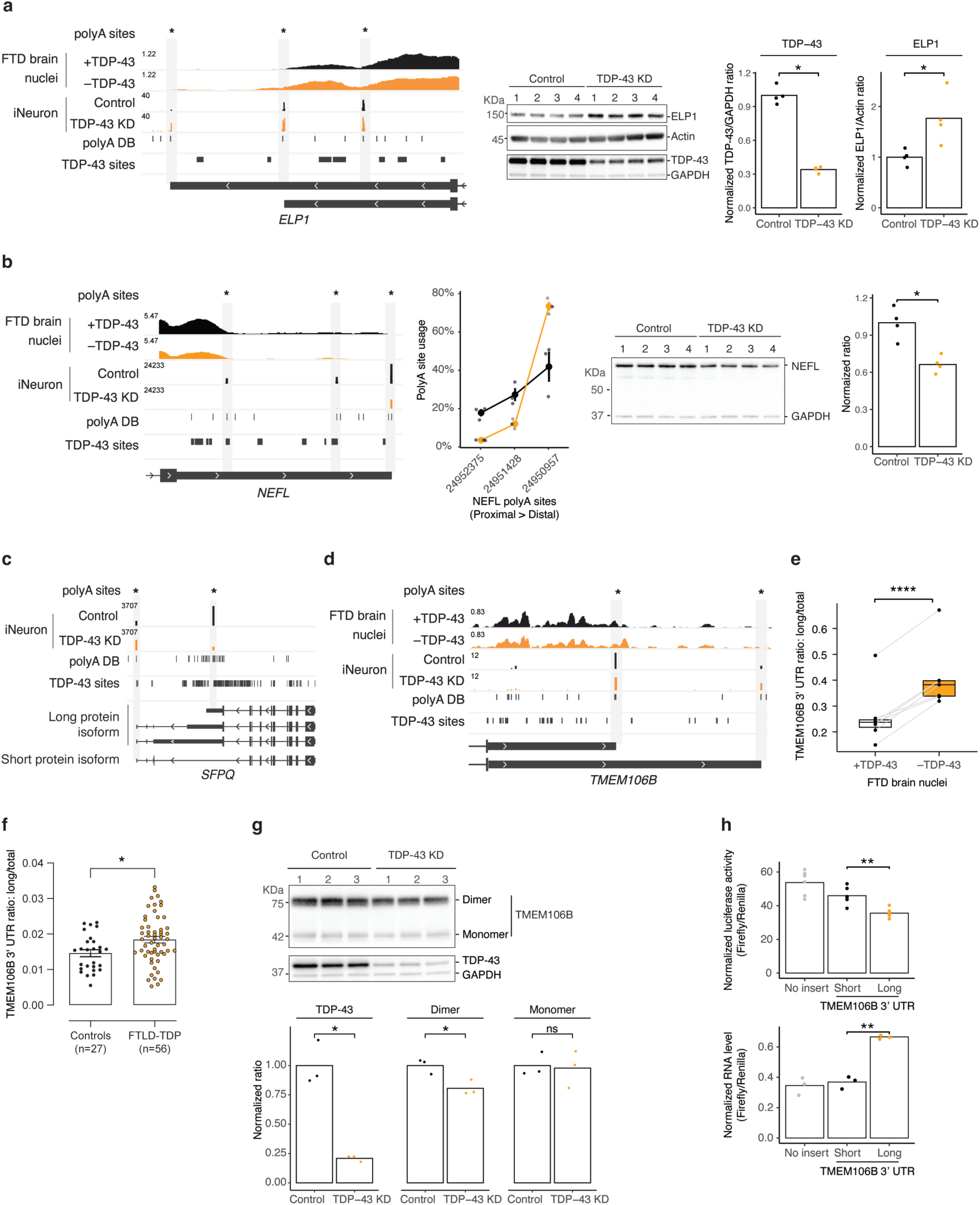
Loss of TDP-43 causes alternative polyadenylation that alters protein levels of disease-associated genes. **a**, TDP-43 knockdown (KD) lengthened the 3’ UTR of *ELP1* (**left panel**) and increased ELP1 protein levels (**middle and right panels**). **b**, TDP-43 KD down-regulates the usage of proximal polyA site usage of *NEFL*, leading to increased usage of the most distal polyA site (**left two panels**) and reduced protein levels (**right two panels**). **c**, TDP-43 KD shifted the polyA site usage from proximal to distal in *SFPQ*. The gene structure shows that the top three RNA isoforms produce a long protein isoform, and the bottom RNA isoform produces a short protein isoform without NLS. **d**, TDP-43 KD lengthened the 3’ UTR of *TMEM106B*. **e**, Targeted APA analysis of *TMEM106B* using RNA-seq from FTD/ALS brain samples. **f**, Level of the long *TMEM106B* 3’ UTR is significantly increased in the frontal cortices of FTLD-TDP patients compared to healthy controls. *P*-values were calculated by two-tailed Mann-Whitney test. **g**, WB shows reduced TMEM106B dimer levels after 12-day TDP-43 KD. **h**, Presence of the long *TMEM106B* 3’ UTR reduces translation efficiency. Bar plots show that longer *TMEM106B* 3’ UTR led to lower luciferase activity (**top panel**) and higher RNA levels (**bottom panel**). No insert, a dual-luciferase reporter without *TMEM106B* 3’ UTRs; short, a dual-luciferase reporter with the short *TMEM106B* 3’ UTR; long, a dual-luciferase reporter with the long *TMEM106B* 3’ UTR. Unless stated otherwise, *p*-values were calculated by Student’s *t* test; ns (not significant), *p* > 0.05; *, *p* ≤ 0.05; **, *p* ≤ 0.01; ***, *p* ≤ 0.001; ****, *p* ≤ 0.0001.

Previous studies found that TDP-43 directly binds to the 3’ UTR of *NEFL* to stabilize its RNA levels^40^. *NEFL* encodes neurofilament light chain (NF-L), which has emerged as a sensitive prognostic biomarker for diverse neurodegenerative diseases^41^, including FTD/ALS^42^. We observed three major polyA sites in *NEFL*, with the most distal one being the most frequently used (**Fig. 3b, left panel**). TDP-43 knockdown not only reduced *NEFL* RNA levels (**Fig. S3e**), consistent with previous findings^43^, but also further shifted the apparent polyA site usage from proximal to distal (**Fig. 3b, left and mid panels; Fig. S3f**) and reduced NF-L protein levels (**Fig. 3b, right panel**). These observations suggest that together with *NEFL* 3’ UTR-targeting microRNAs^44^, loss of TDP-43 induced APA changes might further reduce NF-L levels. A direct role of TDP-43 in regulating NF-L levels adds complexity to the use of this biomarker in TDP-43 proteinopathies.

TDP-43 knockdown also shifted polyA site usage from proximal to distal in another FTD/ALS-linked gene, *SFPQ* (**Fig. 3c**; **Fig. S3g**). *SFPQ* encodes a ubiquitously expressed RNA-binding protein that plays key roles in RNA metabolism^45,46^. *SFPQ* encodes two protein isoforms; the shorter isoform uses a stop codon downstream of the proximal polyA site (**Fig. 3c**) and lacks a nuclear localization signal^45^. Intriguingly, depletion of SFPQ from the nucleus has been observed in sporadic ALS spinal cord^47^, and its interaction with FUS, another FTD/ALS risk gene, is impaired in neuronal nuclei in FTLD-TDP^48,49^. Thus, loss of TDP-43 promotes the usage of *SFPQ*’s distal polyA site (**Fig. 3c**), which might upregulate the shorter NLS-lacking SFPQ isoform and contribute to its nuclear depletion in disease.

The sensitivity of 3’ end-seq further revealed that TDP-43 knockdown lengthened *TMEM106B*’s 3’ UTR (**Fig. 3d**), which we also confirmed by qRT-PCR (**Fig. S3h**). *TMEM106B* is a top genetic risk factor that emerged in a genome-wide association study for FTLD-TDP^50^. C-terminal fragments of TMEM106B were recently found to form amyloid fibrils in brains of older individuals and patients with neurodegenerative disorders^51–54^. By targeted analysis of RNA-seq read coverage across the *TMEM106B* 3’ UTR, we confirmed that loss of TDP-43 is associated with a longer *TMEM106B* 3’ UTR in FTD/ALS postmortem brain samples (**Fig. 3e**). We also analyzed the *TMEM106B* 3’UTR in a series of 83 frontal cortex brain samples from the Mayo Clinic Brain Bank using qRT-PCR and found a significant increase in longer *TMEM106B* 3’ UTRs in the frontal cortices of patients with FTLD-TDP compared with healthy controls (**Fig. 3f**), indicating that the increased 3’ UTR length of *TMEM106B* might be functionally relevant in FTD/ALS.

Because TMEM106B protein levels are altered in disease^50,55^, we next asked whether TDP-43 knockdown affected TMEM106B protein levels. Using a condition that has been shown previously to preserve TMEM106B dimers on SDS-PAGE^55^ (**see also Fig. S3i**), we found that TDP-43 knockdown did not affect levels of TMEM106B monomers (∼42 KDa) but reduced dimer levels (**Fig. 3g**). A recent proteomics study also detected decreased TMEM106B levels caused by TDP-43 knockdown^12^. TDP-43 knockdown only modestly reduced *TMEM106B* RNA levels (**Fig. S3j**), suggesting that the longer 3’ UTR might affect TMEM106B protein levels via translation. To test this hypothesis, we cloned short and long versions of the *TMEM106B* 3’ UTR into a dual-luciferase reporter. We mutated the proximal polyA site in the long 3’ UTR-containing reporter to prevent its usage and confirmed the production of the intended long 3’ UTR (**Fig. S3k**). The long 3’ UTR-containing reporter had significantly lower luciferase activity and higher RNA levels than the short 3’ UTR-containing reporter (**Fig. 3h**), providing evidence that the loss of TDP-43-induced longer *TMEM106B* 3’ UTR reduces translation efficiency. But how could this reduce the formation of TMEM106B dimers? *TMEM106B* APA might impact dimer levels by influencing the subcellular localization of *TMEM106B* mRNA and its ability as a scaffold for protein-protein interactions, as has been shown for other APA events^17^. Recent studies discovered an *Alu* element insertion in the *TMEM106B* 3’ UTR, in perfect linkage with the top FTLD-TDP risk allele at this locus^56–58^. Given the proximity of these variants to *TMEM106B*’s distal polyA site, future work will explore if and how these disease-associated genetic variants impact APA.

In the present study, we applied 3’ end-seq to comprehensively map neuronal polyadenylation on a transcriptomic scale with base-pair resolution and discovered that TDP-43 dysfunction causes widespread APA changes, some of which are in genes critical for neuronal function and in disease-associated genes. These changes are highlighted by coupled cryptic splicing and premature polyadenylation in *ARHGAP32* (**Fig. 2g**), changes observed in multiple subunits of the elongator complex (**Fig. 3a, S3c, S3d**), and 3’ UTR lengthening in *NEFL*, *SFPQ*, and *TMEM106B* (**Fig. 3b-d**). This study is accompanied by two independent manuscripts also presenting widespread APA changes associated with TDP-43 dysfunction in FTD/ALS (Fratta and colleagues^35^; La Spada and colleagues^29^). Despite using different approaches, each of the three studies observed common APA changes (**Fig. 2g**, **S2d-f**, **S2k-m**), underscoring the importance of APA changes during neurodegeneration.

Our application of 3’ end-seq complements and extends RNA-seq based APA analysis because it enables *de novo* identification of polyadenylation events with base-pair resolution, revealing complex and highly sensitive APA changes (e.g., *NEFL*, *ELP3*, and *TMEM106B*). Together, the findings of these three studies provide evidence that TDP-43 pathology contributes to disease pathogenesis through not only cryptic splicing but now also changes in alternative polyadenylation.

## List of supplementary tables

**Table S1: Alternative polyadenylation changes detected in Liu et al. 2019**

**Table S2: Significant APA changes in 3’ UTRs upon TDP-43 knockdown detected by RNA-seq**

**Table S3: Significant APA changes upon TDP-43 knockdown detected by 3’ end-seq**

**Table S4: Curated list of FTD and ALS related genes**

**Table S5: Cryptic polyA sites activated by TDP-43 knockdown**

**Table S6: List of primers used in qRT-PCR**

**Table S7: Summary of patient data**

---

**Figure S1.**
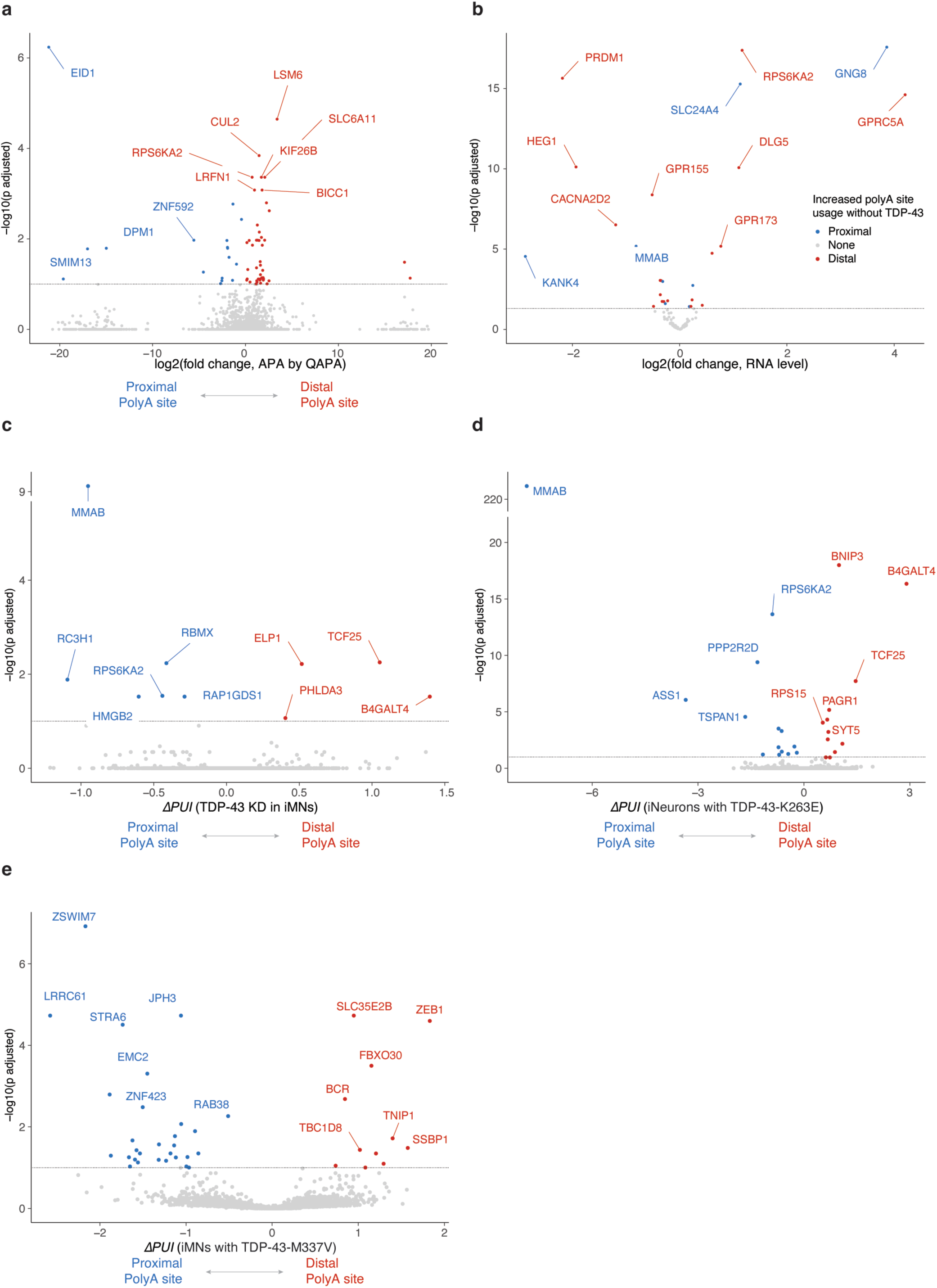
Loss of TDP-43 leads to widespread alternative polyadenylation changes. **a**, Volcano plot shows that QAPA, another popular APA analysis program, also uncovered that loss of TDP-43 is associated with APA changes. The log2(fold change) represents the difference of distal polyA site usage between TDP-43 negative nuclei and TDP-43 positive nuclei. Adjusted p-values were calculated by DEXSeq. **b**, Volcano plot shows that genes have both APA changes and RNA level changes. Only genes with significant APA changes, detected by APAlyzer or QAPA, are plotted with RNA level change on the *x*-axis and the corresponding adjusted *p* value on the *y*-axis. Differential RNA expression was analyzed using DESeq2. **c-e**, Volcano plots show that APAlyzer-identified APA changes upon TPD-43 knockdown (KD) in iMNs (**c**), in iNeurons carrying K263E mutation in TDP-43 (**d**), and in iMNs carrying M337V mutation in TDP-43 (**e**). Details, as in Fig. 1e.

**Figure S2.**
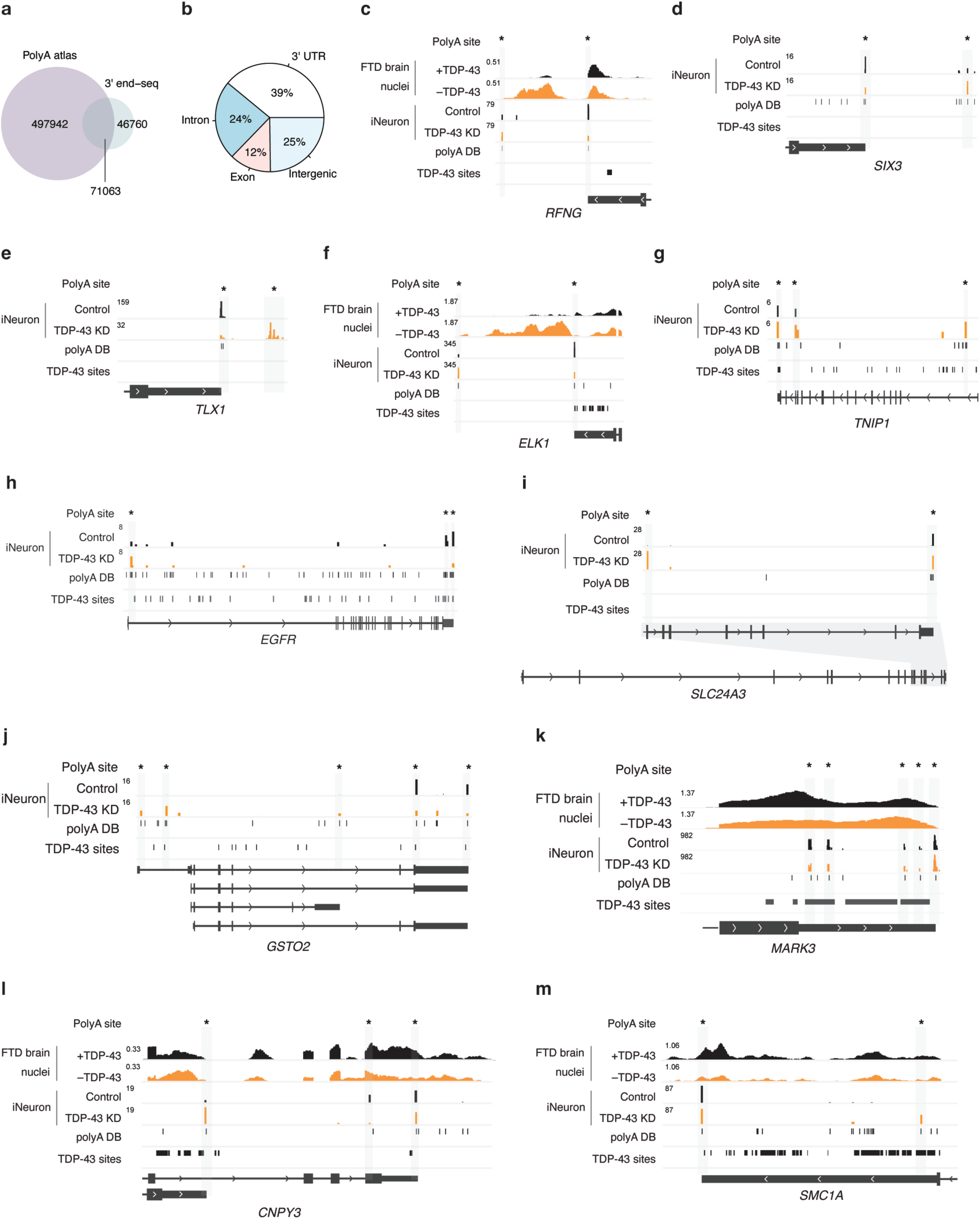
High-resolution mapping of TDP-43 dependent alternative polyadenylation using 3’ end-seq. **a**, Venn Diagram illustrates that 3’ end-seq captured annotated and novel polyA sites. Reads from both control and TDP-43 knockdown (KD) samples were used. **b**, Pie chart shows the distribution of 3’ end-seq identified polyA sites. Note that “intergenic” represents polyA sites not associated with annotated 3’ UTRs, exons, or introns. **c-f**, Examples of the use of an unannotated distal polyA site upon TDP-43 KD. **g-j**, Examples of premature polyadenylation upon TDP-43 KD. **k**, Example of complex usage change of multiple polyA sites upon TDP-43 KD. **l**. Example of increased proximal polyA usage upon TDP-43 KD. **m**, Example of 3’ UTR shortening upon TDP-43 KD. Details, as in Fig. 2g.

**Figure S3.**
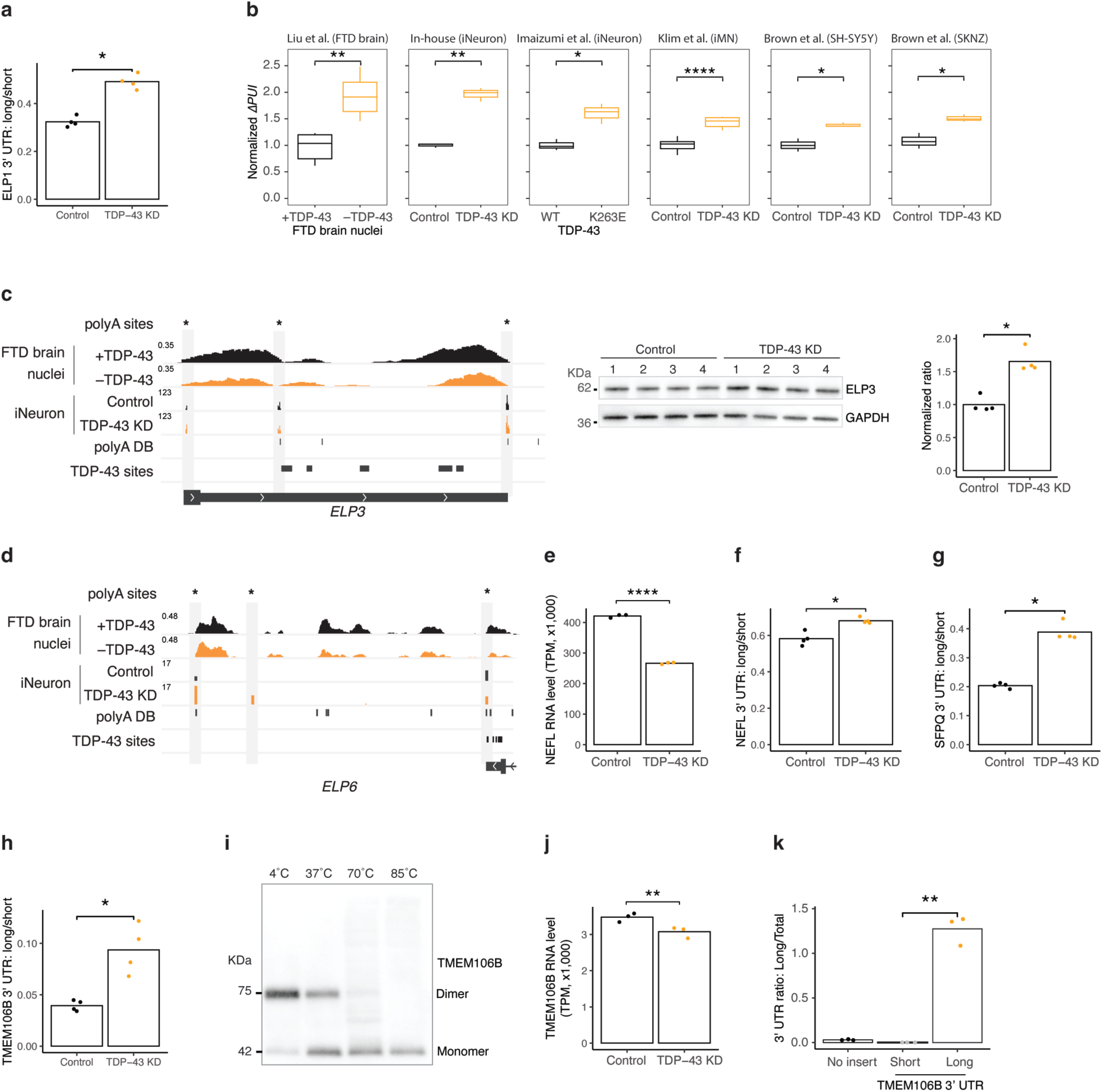
Loss of TDP-43 causes alternative polyadenylation that alters protein levels of disease-associated genes. **a**, Bar plots show that TDP-43 knockdown (KD) increased the level of the long *ELP1* 3’ UTR, confirmed by qRT-PCR. **b**, Bar plots show that higher level of the long *ELP1* 3’ UTR is present across diverse datasets in which TDP-43 was either knocked down or carrying a pathogenic mutation. Normalized *ΔPUI* was calculated as followed: normalized read coverage in the long 3’ UTR region was divided by normalized read coverage in the common 3’ UTR region and the resulting ratio was normalized to the condition that TDP-43 was functional. **c**, TDP-43 KD caused complex APA changes in the 3’ UTR of *ELP3* (**left panel**), increasing its protein levels (**middle and right panels**). **d**, TDP-43 KD lengthened the 3’ UTR of *ELP6*. **e**, Bar plots show that TDP-43 KD reduced *NEFL* RNA levels. The adjusted *p* value was calculated in DEseq2. **f-h**, Bar plots show that TDP-43 KD increased levels of long 3’ UTRs of *NEFL* (**f**), *SFPQ* (**g**), and TMEM106B (**h**), confirmed by qRT-PCR. **i**, WB shows that as the temperature increases, a ∼75 KDa band collapses to a 42 KDa band. Cell lysates were incubated at respective temperatures for 10 min before electrophoresis. **j**, Bar plots show that TDP-43 KD modestly reduced *TMEM106B* RNA levels. The adjusted *p* value was calculated in DEseq2. **k**, Bar plots show that the reporter with the long *TMEM106B* 3’ UTR produced the long 3’ UTR, as designed. RNA levels were measured by qRT-PCR. Unless stated otherwise, *p*-values were calculated by Student’s *t* test. ns (not significant), *p* > 0.05; *, *p* ≤ 0.05; **, *p* ≤ 0.01; ***, *p* ≤ 0.001; ****, *p* ≤ 0.0001.

## Acknowledgments

We thank members of the Gitler lab and the Petrucelli lab for helpful discussions and comments on the manuscript. We thank Abigail Song and Maylin Fu for help with experiments. We thank Ziwei Chen for helping with Aparent2. Y.Z. is supported by a postdoctoral scholar award from The Phil and Penny Knight Initiative for Brain Resilience at the Wu Tsai Neurosciences Institute, Stanford University, and a fellowship grant from the Larry L. Hillblom foundation. T.A. is supported by NIH (2T32AG047126-06A1) and a fellowship from the Takeda Science Foundation. C.G. is supported by Milton Safenowitz Postdoctoral Fellowship Program. Work in A.D.G. is supported by NIH (grants R35NS097263, U54NS123743, R01AG064690) and Target ALS. A.D.G. is a Chan Zuckerberg Biohub – San Francisco Investigator. Work in L.P. is supported by NIH (U54NS123743, R35NS097273, P01NS084974) and Target ALS. Work in M.P. is supported by NIH (U54NS123743, RF1NS120992) and Target ALS.

## Methods

### Cell culture

HEK293T cells were maintained in DMEM (1x) + GlutaMax-I media (Gibco, 10564011) with 10% FBS (Gibco, 16000044) and 100 U/mL Penicillin-Streptomycin (Gibco, 15140122).

### Stem cell maintenance and differentiation into iNeurons

Human embryonic stem cells (hESCs; H1) were maintained in mTeSR1 plus media (StemCell Technologies, 100-0276) on Matrigel (Corning, 354230). hESCs were fed every two days and split every 4–7 days using ReLeSR (StemCell Technologies, 100-0483) according to the manufacturer’s instructions. The differentiation of hESCs to neurons by forcing *NGN2* over-expression was carried out as previously described^60^. In brief, cells were transduced with a Tet-On induction system to drive the expression of the transcription factor *NGN2*. Cells were dissociated on day 3 of differentiation and replated on Matrigel-coated tissue culture plates in Neurobasal Medium (Thermo Fisher, 21103049) containing neurotrophic factors, BDNF and GDNF (R&D Systems).

### TDP-43 knockdown in iNeurons

After seven days of being cultured for differentiation, iNeurons were transduced with lentivirus expressing scramble shRNA or TDP-43 shRNA, cultured for additional seven days or 12 days, and then collected for downstream analyses. The knock-down efficiency was assessed by Western blotting.

### Immunoblotting

After seven days of TDP-43 knockdown, cells were lysed at 4 °C for 15 min in ice-cold RIPA buffer (Sigma-Aldrich R0278) supplemented with a protease inhibitor cocktail (Thermo Fisher 78429) and phosphatase inhibitor (Thermo Fisher 78426). After pelleting lysates at 20,000xg on a table-top centrifuge for 15 min at 4 °C, the supernatant was used for bicinchoninic acid (Invitrogen 23225) assays to determine protein concentrations. Unless stated otherwise, 5-10 ug of protein lysates of each sample was denatured for 10 min at 70 °C in LDS sample buffer (Invitrogen NP0008) containing 2.5% 2-mercaptoethanol (Sigma-Aldrich). These samples were loaded onto 4–12% Bis–Tris Mini gels (Thermo Fisher NP0335BOX) for gel electrophoresis and then transferred onto 0.45-μm nitrocellulose membranes (Bio-Rad 162-0115) using semi-dry transfer method (Bio-Rad Trans-Blot Turbo Transfer System, 1704150) or at 100 V for 2 h at 4 °C using the wet transfer method (Bio-Rad Mini Trans-Blot Electrophoretic Cell 170-3930). Membranes were blocked in EveryBlot Blocking Buffer (Bio-Rad 12010020) or 5% non-fat dry milk in TBST for 1 h then incubated overnight at 4°C in blocking buffer containing antibodies against TMEM106B (1:500, Cell Signaling Technology 93334), TDP-43 (1:1,500, Proteintech 10782-2-AP), ELP1 (1:500, Cell Signaling Technology 5071S), ELP3 (1:1000, Proteintech 17016-1-AP), NEFL (1:1000, Thermo Fisher MA5-14981), GAPDH (1:2,000, Sigma-Aldrich G8795), or GAPDH (D16H11) XP (1:1000, Cell Signaling Technology 8884S). Membranes were subsequently incubated in blocking buffer containing horseradish peroxidase (HRP)-conjugated anti-mouse IgG (H+L) (1:5,000, Fisher 62-6520) or HRP-conjugated anti-rabbit IgG (H+L) (1:5,000, Life Technologies 31462) for 1 h. Amersham ECL Prime kit (Cytiva RPN2232) or SuperSignal™ West Femto Maximum Sensitivity Substrate (Thermo Fisher 34094) was used to develop blots and imaged using ChemiDox XRS+ System (Bio-Rad). The intensity of bands was quantified using Fiji, and then normalized to the corresponding controls.

### Immunoblotting of TMEM106B

After 12 days of TDP-43 knockdown, cells were lysed and normalized as described above. To detect the impact of TDP-43 knockdown on the dimer level of TMEM106B, normalized cell lysates were mixed with 2x laemmli buffer (Bio-Rad) containing 5% 2-mercaptoethanol (Sigma-Aldrich) and loaded directly onto 4-20% Tris-Glycine mini gels (Thermo Fisher XP04205BOX) for gel electrophoresis on ice. To detect the temperature sensitivity of TMEM106B dimer, cell lysates were incubated at 4°C, 37°C, 70°C, and 85°C for 10 min and then loaded onto a 4-20% Tris-Glycine mini gel for gel electrophoresis on ice. After electrophoresis, samples were transferred onto 0.45-μm PVDF membranes (Bio-Rad 162-0115) at 250 mA for 2 h using the wet transfer method (Bio-Rad Mini Trans-Blot Electrophoretic Cell 170-3930). Membranes were blocked in 5% non-fat dry milk in TBST for 1 h and then incubated overnight at 4°C in blocking buffer containing the antibody against TMEM106B (1:500, Cell Signaling Technology 93334). The secondary antibody, imaging, and quantitation are the same as described above.

### Total RNA extraction from iNeurons

Total RNA was extracted using Trizol according to the manufacturer’s instructions. Total RNA was then treated with Turbo DNase (Thermo Fisher AM2238) and cleaned by Zymo’s clean and concentration columns. The quality of RNA was examined by on a High Sensitivity RNA ScreenTape (Agilent, Tapestation).

### qRT-PCR from iNeurons

Total RNA (500 ng) was reverse transcribed to cDNA using the PrimeScript™ RT Reagent Kit with gDNA Eraser (Takara, RR047A). qPCR was carried out using PowerUp™ SYBR™ Green Master Mix kit and detected by QuantStudio3 system (Thermo Fisher). Primers are listed in **Table S6**.

### Total RNA sequencing

Total RNAs from scramble shRNA or TDP-43 shRNA treated cells were used to construct RNA-sequencing libraries using the SMARTer Stranded Total RNA-Seq Kit v2 - Pico Input Mammalian kit (Takara 634411), according to the manufacturer’s instructions. The resulting libraries were quantitated, pooled, and sequenced on a Nextseq 500 machine using the 150-cycle high output kit in a 75bp paired-end mode (Illumina).

### Gene expression analysis

Adaptors in FASTQ files were trimmed using fastp. The adapter trimmed FASTQ files were used for differential gene expression analysis using Salmon and DESeq2.

### Splicing analysis

The adapter-trimmed FASTQ files were mapped to human genome (hg38) following ENCODE recommended settings using STAR. The unique-mapping, properly paired reads were then used for splicing analysis using leafcutter. Cryptic splicing events were called by leafcutter.

### APA analysis using RNA-seq data

Both in-house RNA-seq and publicly available datasets were used for the analysis. The publicly available datasets were downloaded from GEO or SRA (GSE126542, GSE121569, GSE147544, GSE196144, and ERP126666). The adapter trimmed FASTQ files were mapped to human genome (hg38) following ENCODE recommended settings using STAR. The unique-mapping, properly paired reads were then used for APA detection by either APAlyzer or QAPA and then differential analysis by DEXseq. APAlyzer uses polyA DB3 (https://exon.apps.wistar.org/PolyA_DB/) and QAPA uses PolyASite 2.0 (https://polyasite.unibas.ch/) as polyA site reference for detecting APA events.

### 3’end-seq

To comprehensively map polyadenylation and quantify alternative polyadenylation, three control samples and three TDP-43 knockdown samples were used for 3’ end-seq. For each sample, 500 ng of total RNA was used to construct the 3’ end-seq library using Quantseq REV kit from Lexogen, according to the manufacturer’s instructions. The resulting 3’ end-seq libraries were quantified by Qubit, checked for library sizes on a D1000 high sensitivity chip (Agilent Technologies, Tapestation), pooled, and sequenced with on a Nextseq 500 or Nextseq 2000 machine using the 150-cycle high output kit in a 75bp paired-end mode (Illumina).

### APA analysis using 3’ end-seq data

The sequenced 3’ end-seq libraries were adapter-trimmed, and quality filtered using bbduk. The filtered reads were mapped to human genome (hg38) using STAR and then extracted for unique-mapping, properly paired reads using Samtools. If a read was mapped to a region that is immediately upstream of six consecutive As or of a 10-bp region with at least 60% of As, it was considered as a mis-primed read and removed. The resulting filtered reads were analyzed using a modified version of LAPA^61^ that used the default setting, except requiring a replication rate cutoff at 0.75 and can identify polyA site-containing reads mapped to the reverse strand. The change in the polyA site usage was considered significant if the usage difference between two conditions was >10% with adjusted p value < 0.05. PolyA sites were considered cryptic if their usage was ≤5% under the control condition but ≥10% under the TDP-43 knockdown condition and their usage increase was ≥10%. To identify TDP-43 dependent APA events, genes were considered only if they had at least two identified polyA sites and if they had more than two polyA sites, they were filtered for two polyA sites with the two largest usage changes and used for plotting (**Table S3**). Cryptic polyA sites further include polyA sites that became activated upon TDP-43 knockdown and are located downstream of a gene’s annotated 3’ end, which is extracted from a curated gene annotation database. Cryptic splicing events detected in RNA-seq data and 3’ end-seq data were used to search for coordinated cryptic polyadenylation events.

### Calculation of polyA site score

For each polyA site identified in 3’ end-seq libraries, a 205 bp sequence centered at site was extracted and used to calculate the polyA site score using Aparent2, which was then converted to log odds ratio.

### Evaluation of *TMEM106B* 3’ UTR length in FTLD-TDP brain samples by qRT-PCR

Our study cohort included a total of 83 postmortem cases classified into two main groups: healthy controls (n=27) and Frontotemporal dementia (FTD) cases (n=56). Summary of patient data is included in **Table S7**. RNA was extracted from the frontal cortices of the healthy or the FTD patients following the manufacturer’s protocol using the RNAeasy Plus Mini Kit (Qiagen) and as previously described^9,11,62^. Up to three cuts of the sample was used for extraction and only the high-quality RNA samples were processed for downstream analysis. RNA concentration was measured by using Nanodrop technologies (Thermo Fisher) and the RNA integrity number (RIN) was evaluated by Agilent 2100 bioanalyzer (Agilent Technologies). Subsequently, 500 ng of the total RNA extracted was reverse transcribed into cDNA using the High-Capacity cDNA Transcription Kit (Applied Biosystems) per the manufacturer’s instructions. cDNA samples, in triplicates, with SYBR GreenER qPCR SuperMix (Invitrogen), were further evaluated for the quantitative real-time PCR (qRT-PCR) on a QuantStudio™ 7 Flex Real-Time PCR System (Applied Biosystems). Relative quantification of the long *TMEM106B* 3’ UTR and total *TMEM106B* levels was determined using the ΔΔCt method and normalized to two endogenous controls, *GAPDH* and *RPLP0*. All the statistical analyses were performed using the GraphPad Prism 10 (GraphPad Software). For comparison of the frontal cortex RNA levels between the healthy and FTD cases, Mann-Whitney test was used. Primers are listed in **Table S6**.

### Construction of luciferase reporters

The short *TMEM106B* 3’ UTR and the long *TMEM106B* 3’ UTR were amplified from human genomic DNA (H1) and then cloned into the pmirGLO Dual-Luciferase Vector (Promega, E1330) using Gibson assembly. The proximal polyA site in the long *TMEM106B* 3’ UTR was then mutated using Gibson assembly. Mutations that disrupt the proximal polyA site were identified using Aparent2. All of the reporters were confirmed to have correct sequences by whole-plasmid sequencing.

### Measurement of luciferase activity

HEK293T cells were plated on a 96-well plate or on a 24-well plate and transfected with the luciferase reporters that have no insert, the short *TMEM106B* 3’ UTR, or the long *TMEM106B* 3’ UTR using Lipofectamine 3000 (Thermo Fisher, L3000001). Two days later, the transfected cells on a 96-well plate were measured for the luciferase activities using Dual-Glo® Luciferase Assay System (Promega, E2920), according to the manufacturer’s instructions; the transfected cells on a 24-well plate were used for qRT-PCR as described above.

### Quantitation and statistical analysis

All quantification and statistical analyses were done in R. Analysis details can be found in figure legends and the main text. Genome tracks were prepared using IGV. All plots were prepared using ggplot2, ggpubr, patchwork, and ggrepel in R.

## Data and code availability

The sequencing data generated in this study will be available at GEO. The data and code supporting the findings of this study are available from the corresponding authors upon reasonable request. The following publicly available data are used in this study: GSE126542, GSE121569, GSE147544, GSE196144, and ERP126666.

